# A cell cycle-dependent GARP-like transcriptional repressor regulates the initiation of differentiation in *Giardia lamblia*

**DOI:** 10.1101/2021.05.27.446072

**Authors:** Han-Wei Shih, Germain C.M. Alas, Alexander R. Paredez

**Affiliations:** Department of Biology, University of Washington, Seattle, Washington 98195

## Abstract

Transcriptional regulation of differentiation is critical for parasitic pathogens to adapt to environmental changes and regulate transmission. In response to encystation stimuli, *Giardia lamblia* cells shift from G1+S phase of the cell cycle to G2+M. By 2-4 hours, cyst wall proteins are upregulated, indicating that key regulatory steps occur within the first four hours of encystation. However, all characterized transcription factors (TFs) in *Giardia* have only been investigated at later time points of encystation. How TFs initiate encystation and link it to the cell cycle remains enigmatic. Here, we systematically screened six putative early upregulated TFs for nuclear localization, established their dynamic expression profiles, and determined their functional role in regulating encystation. We found a critical repressor, GLP4 that increases rapidly after 30 min of encystation stimuli and downregulates encystation specific markers, including Cyst Wall Proteins (CWPs) and enzymes in the cyst N-acetyl galactosamine (GalNAc) pathway. Depletion of GLP4 increases cyst production. Importantly, we observe that G2+M cells exhibit higher levels of CWP1 resulting from the activation of MYB2, a TF previously linked to encystation in *Giardia*. GLP4 upregulation occurs in G1+S cells, suggesting a role in repressing MYB2 and encystation specific genes in the G1+S phase of the cell cycle. Furthermore, we demonstrate that depletion of GLP4 upregulates MYB2 and promotes encystation while overexpression of GLP4 downregulates MYB2 and represses encystation. Together, these results suggest that *Giardia* employs a dose-dependent transcriptional response that involves the cell cycle regulated repressor GLP4 to orchestrate MYB2 and entry into the encystation pathway.

**Importance:** Transition between life cycle stages is a common feature among parasitic pathogens and its regulation must be optimized to balance persistence of infection with transmission. The early transcription factors (TFs) regulating commitment to differentiate are totally unknown in *Giardia*. In this work, we identified GLP4, a previously uncharacterized GARP-like TF, as an early-acting transcriptional repressor that inhibits G1+S cells from entering the encystation pathway. GLP4 is therefore a key regulator controlling the balance between proliferative growth and terminal differentiation into infective cysts.

## Introduction

To cope with environmental change, protozoa from all eukaryotic supergroups differentiate to dormant walled cysts, a process which is commonly called encystation (1). Deposition of a cyst wall to form water-resistant cysts allows parasitic protozoa to persist in fresh water and successfully transmit infection to susceptible hosts via a fecal-oral route. *Giardia lamblia* (synonymous with *Giardia duodenalis* and *Giardia intestinalis*) is a major cause of protozoan-based diarrhea and its encystation pathway plays an important role in pathogen virulence. *Giardia* exhibits a simple biphasic life cycle, consisting of cysts and trophozoites. *Giardia* infection occurs through consuming infectious cysts from contaminated water (2), food, or interpersonal contact (3). Following passage of cysts through the acidic environment of the stomach, vegetative trophozoites excyst in the duodenum and rapidly colonize the small intestine(4). In the small intestine, cell density, lipid starvation, and alkaline pH trigger a portion of trophozoites to initiate encystation (5–8). During the early stage of *Giardia* encystation, major morphological changes occur that include biogenesis of the Golgi-like Encystation Specific Vesicles (ESVs), Encystation Carbohydrate-positive Vesicles (ECVs), cell cycle arrest, and cytoskeleton rearrangement (5, 6, 9). *Giardia’s* cyst wall is composed of 37% cyst wall proteins (CWPs) and 63% carbohydrate filaments which are mainly made up by a [D-GalNAc-β(1-3)-D-GalNAc]_n_ homopolymer (10–12). The expression levels of CWP1-3 are upregulated at 2-4 h post induction of encystation (13–15). The enzymes involved in the synthesis of β(1-3) GalNAc homopolymer, including GlcN 6-P isomerase (G6PI-B), GlcN 6-P N-acetylase (GNPNAT), phosphoacetylglucosamine mutase (PGM), UDP-GlcNAc pyrophosphorylase (UAP), UDP-GlcNAc 4’-epimerase (UAE) are upregulated at 10-18h post induction of encystation (10, 11, 13, 15).

Cell cycle arrest appears to be a common theme in encystation. Studies in *Giardia lamblia* (16), *Acanthamoeba* (17), and *Entamoeba invadens* (18) have all shown that encystation stimuli correlate with the accumulation of G2+M cells in the early stage of encystation. In *Acanthamoeba*, encystation frequency correlates with the proportion of cells arrested at G2(17), suggesting that differentiation may be initiated from G2+M cells. In *Giardia*, aphidicolin-treated trophozoite exhibited more ESVs in response to encystation stimuli. These observations led to a hypothesis that there is also a decision point in *Giardia’s* cell cycle that limits encystation to cells in G2+M (16, 19).

A recent study has shown that a Myeloblastosis domain protein (MYB) transcription factor (TF) is a master regulator of differentiation in *Toxoplasma gondii* (20). MYB TFs also modulate encystation in *Giardia* (21) and *Entamoeba* (22). Proteomic and RNA sequencing studies agree that *Giardia’s* MYB2 TF is a key regulator of encystation (14, 23, 24). In *Giardia*, several TFs have been characterized, including GARP-like protein 1 (GLP1, named for, Golden2, ARR-B, Psr-1 like protein 1) (25), E2 promoter binding factor 1 (E2F1)(26), WRKYGQK domain proteins (WRKY) (27), Paired box proteins 1,2 (PAX1,2)(28, 29), AT rich interaction domain (ARID) protein 1 (30), and MYB2. Thus far, all *Giardia* TFs which have been identified to regulate encystation activate CWPs (21, 25–27, 29, 30). Interestingly, no single repressor has been identified, but promoter analysis studies have suggested the presence of repressor binding sites in CWP2 and G6PI-B (31–33).

The identification of TFs regulating entry into the encystation pathway is critical for understanding how the initiation of differentiation is controlled. In *Giardia*, cell cycle arrest and CWP1 upregulation occur 2-4 h post induction of encystation. Thus far, functional studies of *Giardia’s TFs* have focused on transcriptional regulation 24 h after inducing encystation. The regulatory mechanism of TFs involved in the initiation of encystation and the spatiotemporal dynamic of TF expression at the earliest stage of encystation remain unclear. Here, we addressed these questions by screening putative TFs identified in a previous RNAseq study (14). We determined the expression profiles of 6 TFs upregulated early in encystation, investigating their functional roles by knockdown using CRISPRi and CasRX approaches, which led to the identification and characterization of GLP4, a cell cycle-regulated repressor of encystation. Our findings provide the first evidence of TF dynamics at 30 min to 4 h post induction of encystation. We establish a key molecular link between encystation initiation and GLP4 expression that indicates GLP4 is a cell cycle-dependent repressor that inhibits the entry into encystation by repressing MYB2 expression in G1+S cells.

## Results

### Characteristics of transcription factors during early encystation in *Giardia*

Given that CWP1-3 are canonical encystation indicators in *Giardia (5, 6, 15, 23, 34)*, we first investigated CWP1-3 protein levels during encystation initiation using NanoLuc reporters (35, 36). CWP1-3 protein levels began to increase after 1.5 h and rose exponentially by 4 h after exposure to encystation medium (Fig. 1A). To date, several TFs have been identified with roles in regulating CWP1-2; these include MYB2 (21), E2F1 (26), PAX1-2 (28, 29), ARID1(30) and WRKY (27). Whether these TFs have a role in initiating entry into the encystation pathway has not been established. The increase of CWPs is likely to be signaled through early response TFs, which trigger the activation or repression of encystation specific genes. A previous transcriptomics study identified 6 TF candidates that are upregulated within 4 h of encystation (14). We sought to test whether these TFs function as upstream regulators of encystation. We expressed these 6 proteins as mNeonGreen (mNG)-fusion to verify nuclear localization during early encystation, using MYB2 as a positive control. MYB2 expression has been detected as early as 7 h post induction of encystation in proteomic (23) and RNA sequencing studies (14). We were able to detect nuclear localization of MYB2 at 4 h but not at 0 and 1.5 h (Fig. 1B). In contrast, we observed 6 TFs, including MYB1, GLP3, GLP4, PAX1, ARID2, and E2F1 with nuclear localization at 0 and 1.5 h of encystation stimulus (Fig. 1C; Fig. S1). MYB1 and GLP4 have not been previously studied. We verified that MYB1-mNG and GLP4-mNG localized to the nuclei by co-localization with the fluorescent DNA dye DRAQ5 (Fig. 1C).

**Fig. 1.**
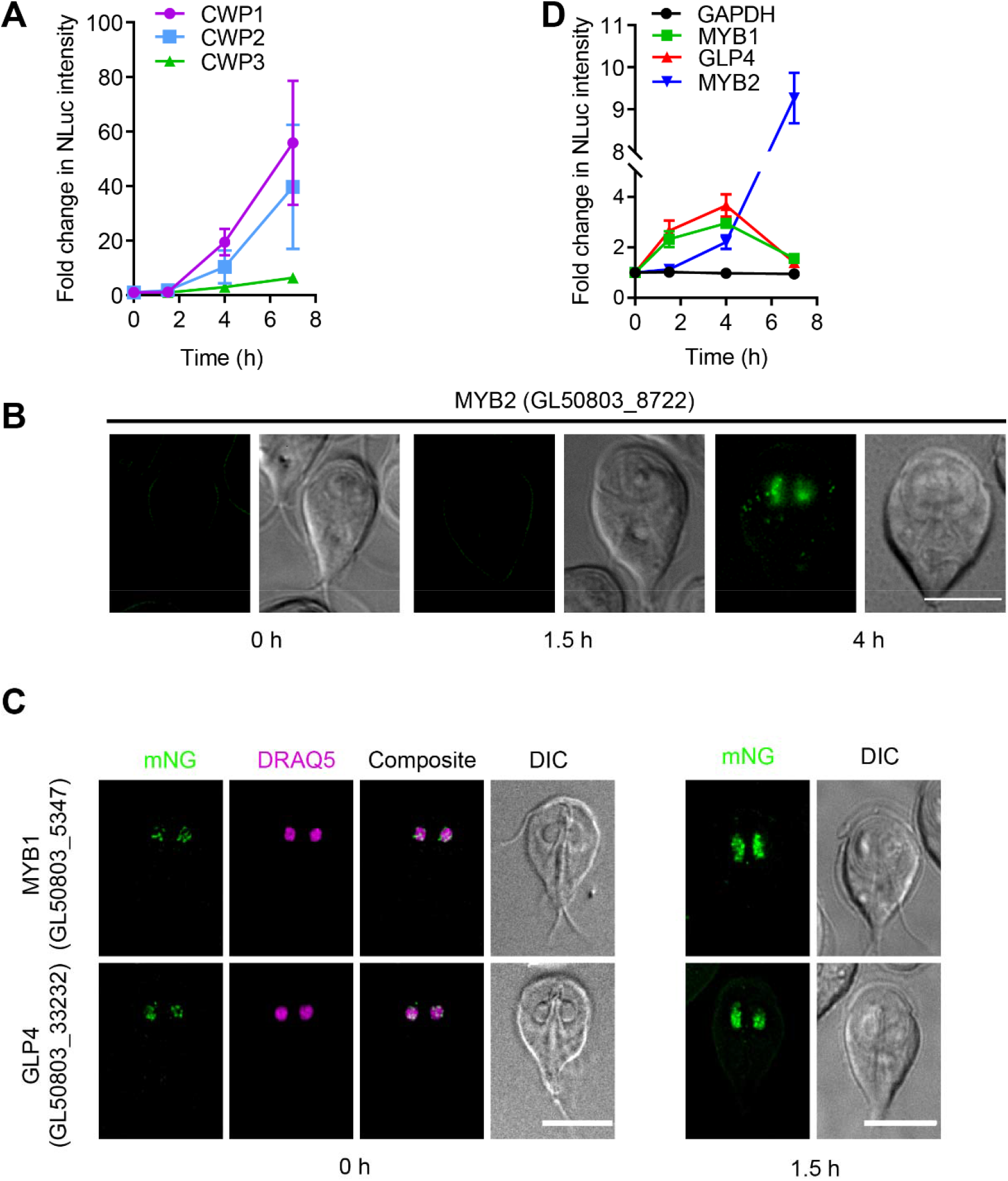
GLP4 and MYB1 are rapidly upregulated TFs. *(A)* Relative NanoLuc (NLuc) luciferase intensity of endogenously tagged CWP1-NLuc, CWP2-NLuc, and CWP3-NLuc after 0, 1.5, and 4 h exposure to encystation medium. The expression level for each time point has three biological replicates. Fold change is normalized to 0 h. Luciferase intensity is measured by plate reader. Each well has 2 × 10^5^ cells. *(B)* Nuclear localizations of MYB2-mNG (GL50803_8722), (C) MYB1-mNG (GL50803_5347) and GLP4-mNG (GL50803_33232) after 0, 1.5, 4, and 7 h exposure to encystation medium. MYB1-mNG and GLP4-mNG were colocalized with nuclear DRAQ5 to show nuclear localization at 0 h post induction of encystation. All images were taken with same exposure. Bars, 10 μm. *(D)* Same quantification as (A) with GLP4-NLuc, MYB1-NLuc, MYB2-NLuc, and GAPDH-NLuc after 0, 1.5, 4, 7h exposure to encystation medium.

We then decided to investigate the expression level of these TFs. GLP4 and MYB1 rose nearly three-fold after 1.5 h of inducing encystation, with GLP4 having the highest level of upregulation (Fig. 1D). Expressions of GLP3, PAX1, ARID2 and E2F1 only modestly increased by 4 h (Fig. S2A). Notably, MYB1 and GLP4 are upregulated before MYB2. This hints that these transcription factors may be early encystation regulators upstream of MYB2 and CWP1.

### Depletion of MYB2 and GLP4 alter cyst production

We next screened for knockdown constructs that would allow us to test the role of the six TFs in regulating encystation. We tested the knockdown efficacy of an established dCas9-based CRISPRi knockdown system (34) and generated a complementary CasRX knockdown system (37) that targets mRNA for degradation (Fig. S3). Based on endogenously-tagged NanoLuc reporter fusions for each of the TFs, we were able to significantly knockdown all six TFs of interest (Fig. S4). Subsequently, we tested the most efficacious knockdown constructs for their ability to alter CWP1 expression.

Previous studies have suggested a critical role for MYB2 in controlling *Giardia* encystation (21, 38); therefore, we investigated the phenotype of a CRISPRi-mediated MYB2 knockdown mutant (Fig. 2A). Depletion of MYB2 reduced CWP1 and CWP2 levels at 4h of encystation (Fig. 2B-C; Fig. S6E) and totally abolished the cyst maturation (Fig. 2D; Fig. S7A), confirming that MYB2 is essential for encystation. Notably, our MYB2 CRISPRi cell line has better performance than a previously reported MYB2 antisense silencing mutant that only partially reduced cyst production (21). E2F1 (Fig. S5A-C) and PAX1 (Fig. S5D-F) have also been reported to regulate CWP levels (26, 29), in agreement, we found that their knockdowns reduced CWP1 levels.

**Fig. 2.**
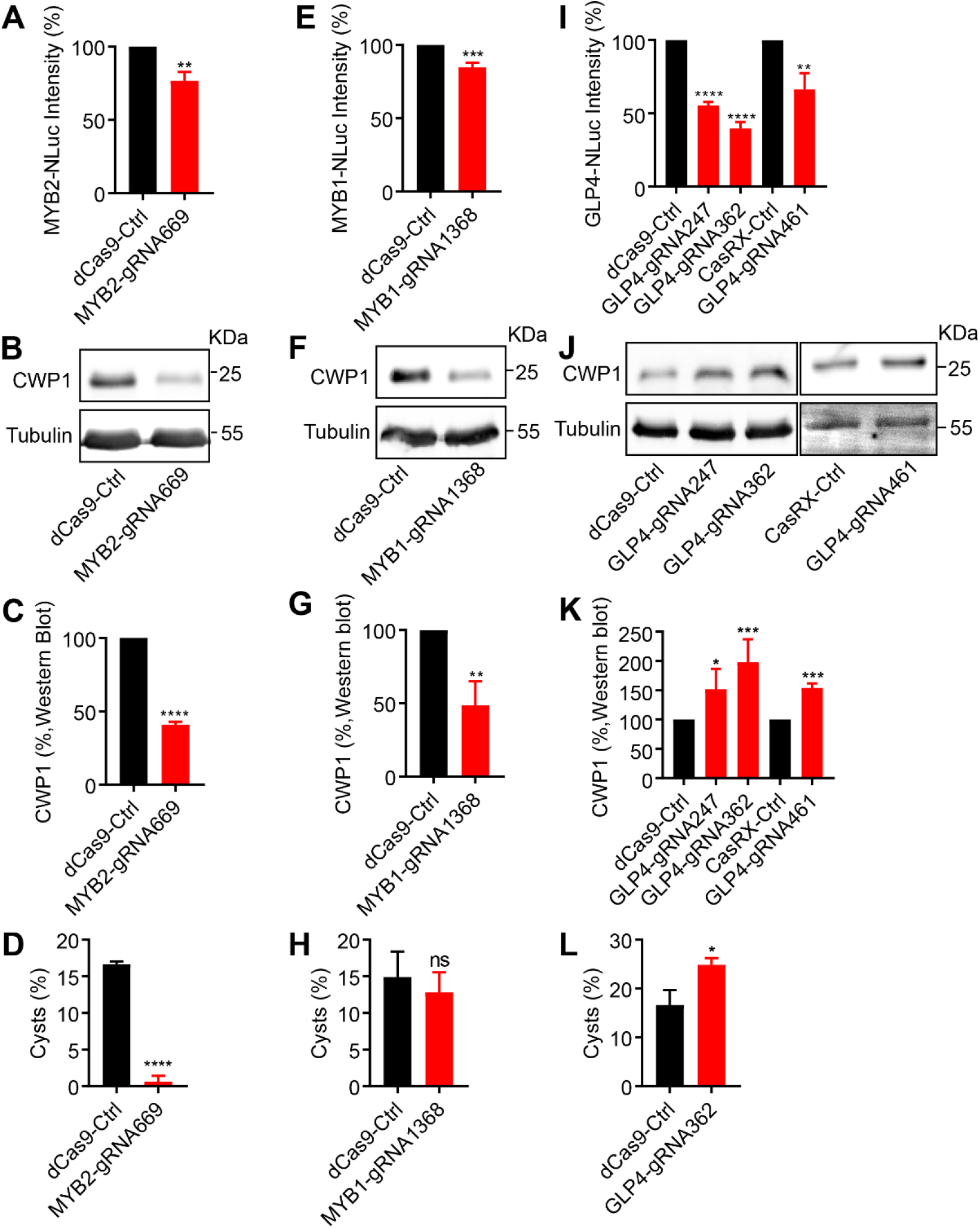
Depletion of GLP4 increases CWP1 expression level and cyst number. *(A,E,I)* Quantification of NanoLuc intensity from CRISPRi or CasRX-mediated knockdowns of (A) MYB2 (GL50803_8722), (E) MYB1 (GL50803_5347), and (I) GLP4 (GL50803_33232). The expression level is normalized to the dCas9 control. *(B,F,J)* Western blots of CWP1 and tubulin from (B) MYB2, (F) MYB1, (J) GLP4 knockdowns at 4h post induction of encystation. *(C,G,K)* Quantification of Western blots of CWP1 and tubulin from (B) MYB2, (F) MYB1, (J) GLP4. The expression level is normalized to tubulin. *(D,H,L)* Quantification of cyst number for (D) MYB2, (H) MYB1, and (L) GLP4 knockdowns after 48 h encystation quantified by hemocytometer. All quantification is from three independent biological replicates. Data are mean ± s.d. (n>100,000 cysts analyzed) Student’s t-test, *p<0.01, **p<0.001, ***p<0.0001, ****p<0.00001, ns= not significant.

The functional role of ARID2, MYB1, and GLP4 in encystation has not been previously established. Knockdown of ARID2 did not alter CWP1 levels (Fig. S5G-I) nor change the portion of cells that enter the encystation pathway (Fig. S6C). Knockdown of MYB1 (Fig. 2E) resulted in lower CWP1 levels (Fig. 2F), lower CWP2 levels (Fig. S6E), and fewer encysting cells at 4 h (Fig. S6D); however, after 48 h the percentage of viable mature cysts (Fig. 2H; Fig. S7B-C) was similar to the control. This suggests that MYB1 might be an upstream regulator of CWP1 but ultimately it is not a major factor in cyst maturation. Interestingly, depletion of GLP4 (Fig. 2I) resulted in a two-fold increase in CWP1 levels (Fig. 2J-K) and an approximately 20% increase in CWP2 levels (Fig. S6E). Notably, GLP4 knockdown resulted in more mature cysts (Fig 2L) without changing cyst viability (Fig. S7D-E). In summary, these results show that GLP4 is a repressor of CWP1 and cyst development and, that while MYB1 is an activator of CWP1, it is not essential for cyst maturation.

### Depletion of GLP4 upregulates the expression of encystation markers

To further characterize the role of GLP4, we investigated whether depletion of GLP4 changes cell growth rate and cell viability. Knockdown of GLP4 did not alter proliferation or viability using the ATP-dependent click beetle green 99 (CBG99) luciferase assay (Fig S8A-B)(39). In addition to the high bile Uppsala encystation method used above, *Giardia* can be encysted with additional methods. These include the two-step method that removes bile and then provides bile levels moderately above the critical concentration for bile acid micelle formation to sequester cholesterol or cells can be starved for cholesterol with lipo-protein deficient media (40). Similar to the high bile Uppsala method, both two-step and lipo-protein deficient encystation media upregulated CWP1 more in the GLP4 mutant background (Fig. S8C), indicating that the GLP4 knockdown phenotype is not restricted to a specific encystation protocol. Considering that GLP4 has an important role in regulating encystation, we examined how its levels change over a fine time course at the induction of encystation. Remarkably, GLP4 levels rapidly increased by 30 min while MYB1 did not appreciably ramp up until after 60 min of encystation (Fig. 3A). GLP4 mutants have consistently higher CWP1 levels during early encystation (Fig. 3B). We analyzed individual cells to determine whether the increase of CWP1 in the GLP4 knockdown lines was due to increased numbers of cells entering the encystation program or resulted from increased levels of CWP1 per cell. We collected encysting cells from different time points of encystation and stained with a CWP1 antibody to quantify encysting cells. We then investigated various phenotypes of CWP1 trafficking in GLP4 mutants. The percentage of encysting cells was 1.5-2 fold higher in GLP4 mutants at 0, 2, 4, and 24 h treatments (Fig. S9; Fig. 3C-D). Furthermore, we analyzed CWP1-positive vesicle size and volume to address whether CWP1 trafficking is altered as a result of increased CWP1 expression in the GLP4 mutant. Our results show that neither vesicle size nor vesicle volume are changed (Fig. 3E-F; Fig. S8D). This indicates that the increased levels of CWP1 are due to more cells entering the encystation pathway. In contrast, overexpression of GLP4 inhibits 50% of CWP1 expression (Fig. S8E-G).

**Fig. 3.**
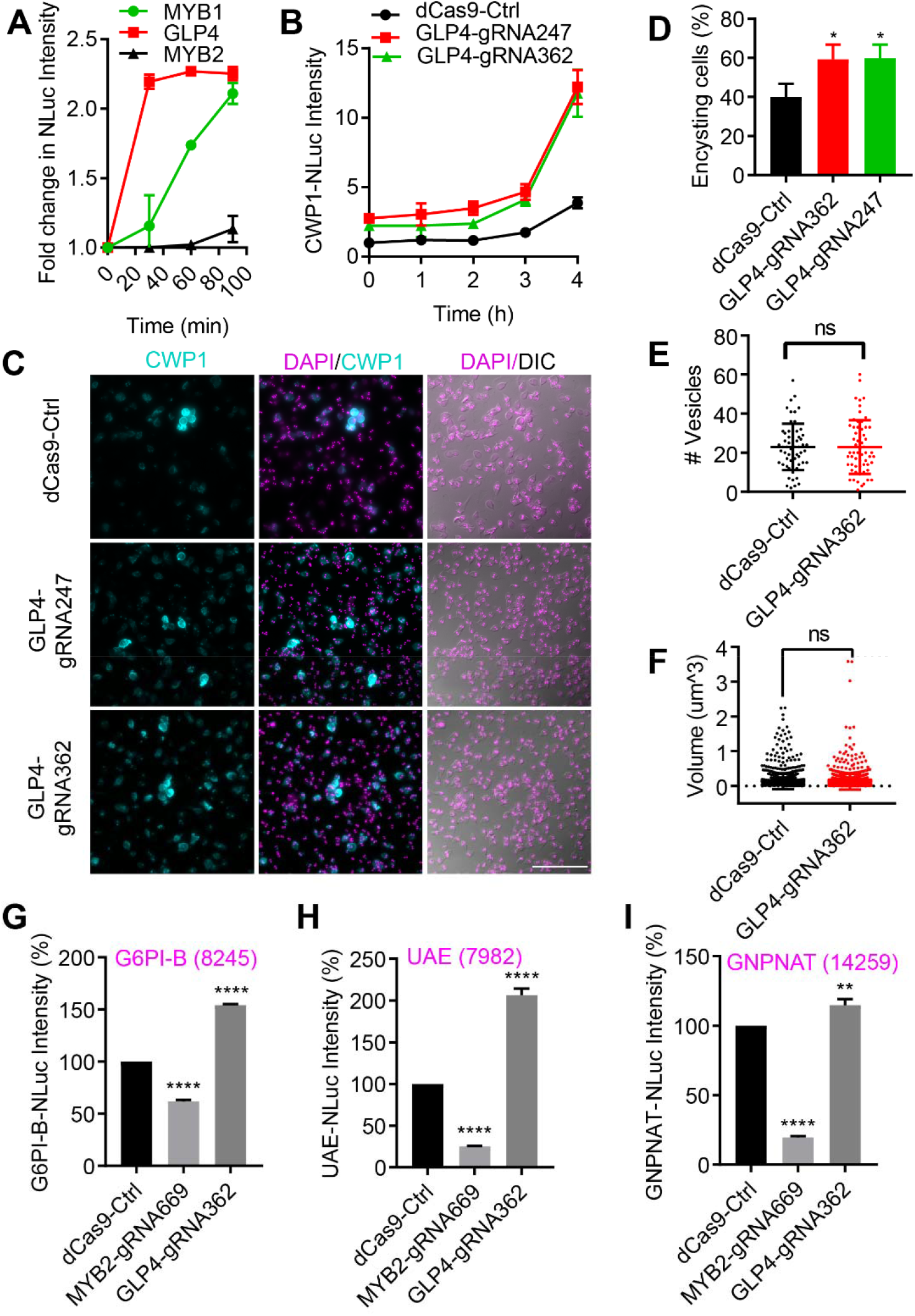
GLP4 downregulates encystation markers. *(A)* Relative NanoLuc intensity of MYB1-NLuc, MYB2-NLuc, and GARP4-NLuc after 0, 0.5, 1 and 2 h exposure to encystation medium (Uppsala medium). The expression level from each time point has three biological replicates. The fold change is normalized to 0 h. *(B)* Relative CWP1-NLuc intensity in GLP4 knockdown cells after 0, 0.5, 1,2,3 and 4 h exposure to encystation medium (Uppsala medium). The expression level from each time point has three biological replicates. The fold change is normalized to 0 h of the dCas9-Ctrl. *(C)* CWP1 and DAPI staining at 24 h of encystation for dCas9-Ctrl, GLP4-gRNA247, and GLP4-gRNA362 cell lines. Bars, 100 μm. *(D)* Quantification of (C). Data are mean ± s.d. from three independent experiments (total cells counted for dCas9-Ctrl n=1830, GLP4-gRNA362 n=1747, GLP4-gRNA247 n=1783) Student’s t-test, *p<0.01. IMARIS-assisted analysis of *(E)* vesicle number and *(F)* vesicle volume in 4 h encysting cells of dCas9-Ctrl, GLP4-gRNA247 and GLP4-gRNA362 cell lines. n.s.= not significant. n=63 cells for dCas9-Ctrl. n= 93 for GLP4-gRNA362. *(G-I)* Relative expression levels of GalNAc enzymes for (G) G6PI-B-NLuc (GL50803_8245), (H) UAE-NLuc (GL50803_7982), and (I) GNPNAT-NLuc (GL50803_14259) in dCas9 control, MYB2-gRNA669, and GLP4-gRNA362 backgrounds. **p<0.001, ****p<0.00001.

Because cyst wall composition includes 67% N-acetyl-D-galactosamine (GalNAc) homopolymers, we sought to analyze whether GLP4 and MYB2 regulate other encystation markers, such as enzymes in the GalNAc pathway (11). *In vivo* transcriptional regulation identified GNPNAT and UAE as highly expressed enzymes in the GalNAc pathway (15). Additionally, MYB2 was shown to modulate G6PI-B (41). Using NanoLuc as a reporter, we analyzed the expression level of G6PI-B, UAE, and GNPNAT at 0, 1.5, 4, 7, 18, and 24 h post induction of encystation. Our data shows that UAE is slightly increased by 1.5 h and G6PI-B began to increase by 4 h (Fig. S2B). At 18 h, UAE, G6PI-B, and GNPNAT levels increased by 150, 20, and 28 fold, respectively (Fig. 2B-C). We next expressed MYB2 and GLP4 CRISPRi guide RNAs individually in G6PI-B, GNPNAT, and UAE NanoLuc cell lines to investigate whether MYB2 and GLP4 modulate their expression. Consistent with our CWP1-2 results, the levels of G6PI-B, UAE, and GNPANT are downregulated in MYB2 knockdown but are upregulated by GLP4 knockdown, suggesting GLP4 is also a repressor of enzymes in the GalNAc pathway (Fig. 3G-I).

### GLP4 is a cell cycle dependent repressor

Encystation-induced G2 arrest has been demonstrated in *Giardia* (42, 43). Indeed, our data is consistent with previous studies that found 1.5 h of encystation is sufficient to cause accumulation of G2+M cells with a corresponding reduction of cells in G1+S (Fig. 4A-B). A previous study showed that *Giardia* trophozoites arrested at G2 with aphidicolin, produced more ESVs after release into encystation medium (42). This study therefore hypothesized that *Giardia* enter the encystation program at G2. However, evidence to correlate CWP1 and MYB2 levels with DNA content during encystation is still lacking. In *Giardia*, nocodazole and aphidicolin treatments or counterflow centrifugal elutriation (CCE) have been used to synchronize trophozoites but there are two technical challenges: (1) Nocodazole and aphidicolin created endoreplication and double-stranded DNA breaks, respectively, and secondary effects resulted in unhealthy cells that would not be ideal for encystation assays (44); (2) the CCE method was performed in 1XHEPES-buffered saline (HBS) and *Giardia* trophozoites are not able to encyst in HBS buffer (44). To more precisely define CWP1 and MYB2 expression levels during encystation at G1 and G2 phases, we analyzed encysting cells with flow cytometry. CWP1 and MYB2 were endogenously tagged with mNG and the cell lines were exposed to encystation stimuli. DNA content was measured by DRAQ5 fluorescence intensity using flow cytometry. A control experiment was performed to verify that the mNG tag does not change CWP1 production in response to encystation stimuli (Fig. S10A-B). We found that cells with higher DNA content had higher CWP1 levels (Fig. S10C). We also used MYB2-mNG to show that G2+M cells have higher MYB2 levels (Fig. S10D), suggesting higher expression of CWP1 was due to higher levels of MYB2. In contrast with the previous conclusion that only G2 cells enter the encystation pathway (42), it is noteworthy that our result indicates that G1 cells produce both CWP1 and MYB2 at 1.5 h post induction of encystation, suggesting *Giardia* enter the encystation program at both G1 and G2 phases instead of G2 only. Given that G2 cells have higher expressions of CWP1 and MYB2, we sought to characterize the expression level of GLP4 in G1 cells. We examined the relationship between GLP4-mNG and the cell cycle at 0 and 1.5 h of encystation. We found that GLP4-mNG fluorescence intensity was increased by 1.5 fold in G1+S cells at 1.5 h post induction of encystation but slightly decreased in cells at G2+M (Fig. 4C), suggesting lower GLP4 levels permit G2+M cells to produce more MYB2 and CWP1 during encystation. Because MYB2 and CWP1 levels are lower in G1+S cells, we questioned if GLP4 might be an upstream repressor of MYB2. We transfected the GLP4 CRISPRi guide RNA into the MYB2-NanoLuc expressing cell line. We found that MYB2 levels rose by 1.2 fold in the GLP4 knockdown cell line at 4h post inducing encystation, which is consistent with MYB2 ultimately regulating CWP1 levels (Fig. 4D). In contrast, overexpression of GLP4 downregulated MYB2 expression by 50% (Fig. 4E-F) and resulted in reduced CWP1 levels (Fig. S8E). These findings suggest MYB2 is downstream of GLP4.

**Fig. 4.**
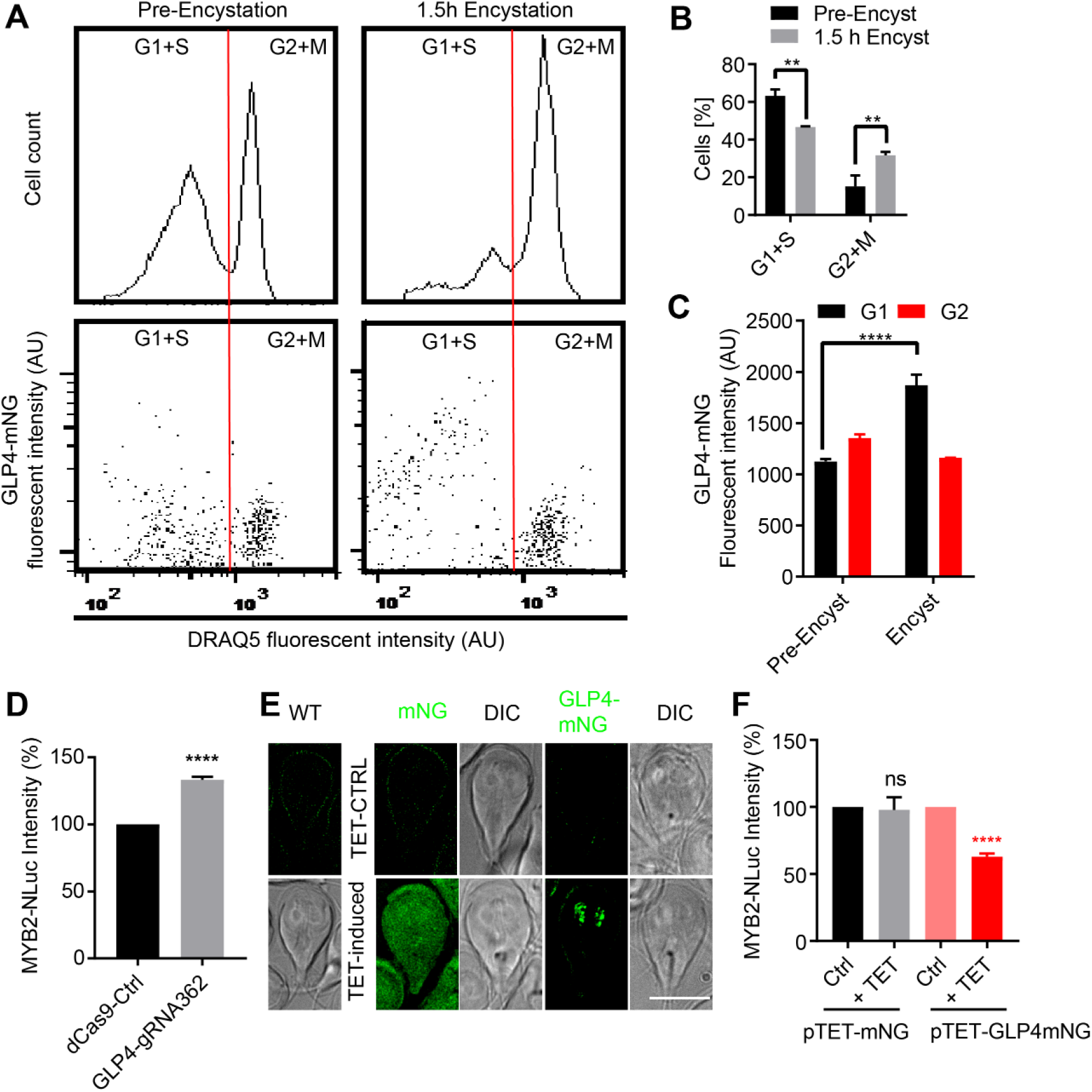
GLP4 is a cell cycle dependent repressor. *(A)* Flow cytometry analysis of DRAQ5 and GLP4-mNG at 0 and 1.5 h of encystation. Red line is the separation point between G1+S and G2+M phases. *(B)* Quantification of G1+S and G2+M cells from 0 and 1.5 h encystation in (A). Data are mean ± s.d. (n=3, 10,000 cells per replicate) Student’s t-test, **p<0.001. *(C)* Quantification of GLP4-mNG fluorescence intensity in G1+S and G2+M cells at 0 and 1.5 h of encystation. Data are mean ± s.d. (n=3) Student’s t-test, ****p<0.0001. *(D)* Relative NanoLuc intensity of MYB2 from dCas9 control and GLP4-gRNA362 cell lines. ****p<0.00001. *(E)* Green fluorescence and DIC images of WT, GLP4-mNG (Int) (integrated and endogenously tagged GLP4-mNG), pTET-mNG (mNG under the tetracycline inducible promoter) and pTET-GLP4-mNG (GLP4-mNG under the tetracycline inducible promoter) in the MYB2-NLuc background with or without tetracycline induction. All images were taken with same exposure. Bars, 10 μm. *(F)* Relative expressions of MYB2-NLuc intensity in mNG and GLP4-mNG overexpression cell lines after 4 h of encystation. Expression levels are normalized to the tetracycline controls at 4 h post induction of encystation. ****p<0.00001. 20 μg/ml tetracycline (final concentration) was added to induce expression for 24 h. ****p<0.00001. n.s.= not significant.

Because E2F1 and PAX1 have been shown to upregulate CWP1-3 and MYB2 (26, 29), we sought to determine whether GLP4 modulates MYB2 through either E2F1 or PAX1 expression levels. Interestingly, knockdown of GLP4 did not alter E2F1 or PAX1 expression levels as measured with NanoLuc reporters (Fig. S10F-G), suggesting E2F1 and PAX1 are not downregulated by GLP4. Thus, MYB2 is a downstream of GLP4 but an upstream regulator of CWP1-2 and enzymes in the GalNAc pathway. In summary, GLP4 has a role in limiting entry into the encystation pathway where MYB2 is repressed in G1+S cells by GLP4, then during G2+M, GLP4 levels drop and MYB2 expression is upregulated to produce more CWPs and enzymes in the GalNAc pathway (Fig. 5).

**Fig. 5.**
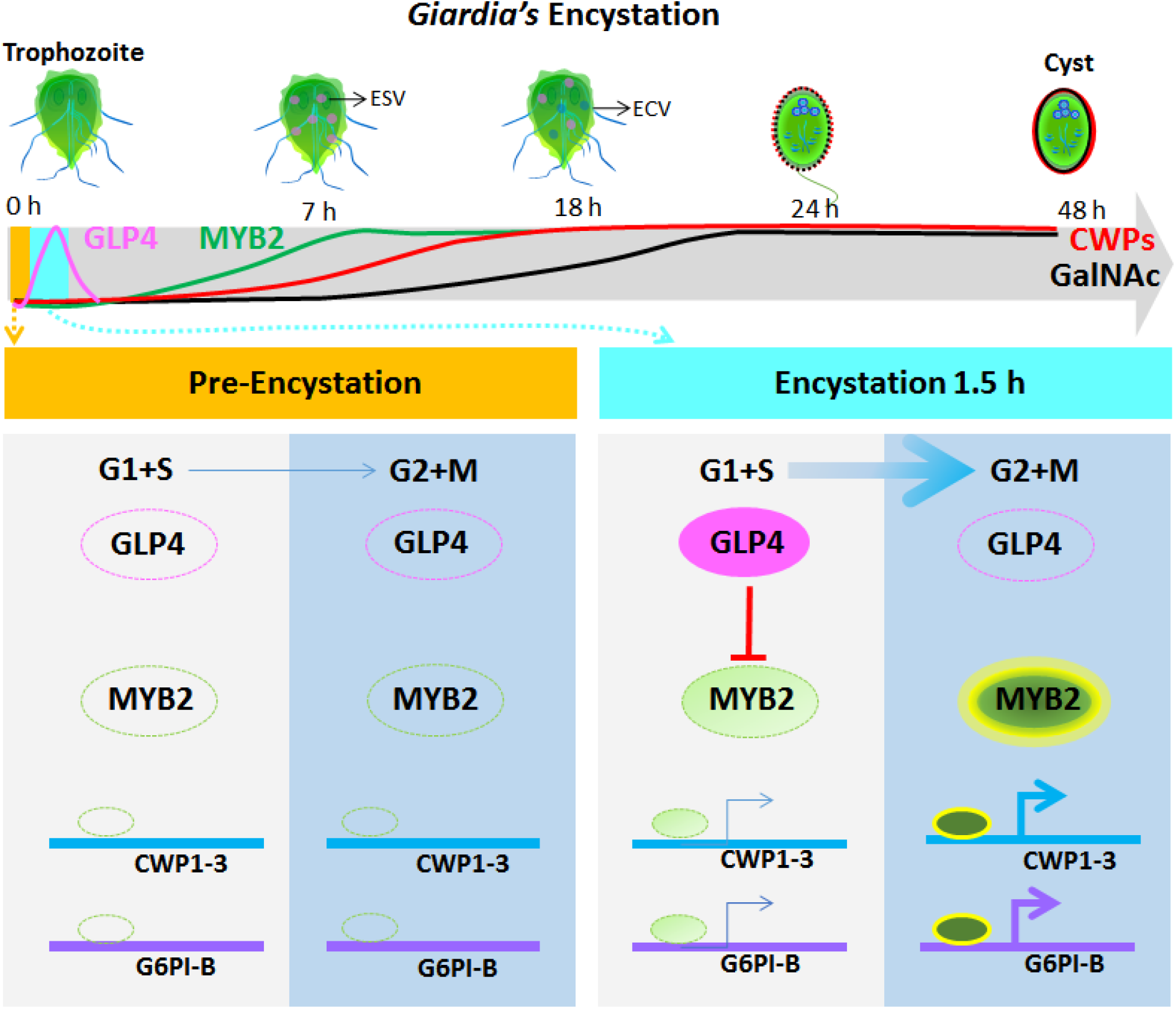
Schematic overview of the dosage dependent transcriptional responses during encystation. ***Top panel*** shows a timeline for encystation that includes 7 h marked by encystation specific vesicles (ESVs, closed pink circle) that traffic CWPs, the 18 h middle stage with ESVs and the encystation carbohydrate-specific vesicles (ECVs, closed blue circle) that traffic the GalNac sugars, the 24 h middle-late stage with cysts that have begun disassembly of their cytoskeletons, and the 48 h late stage with mature cysts. ***Middle panel*** shows the expression profiles of GLP4 (magenta), MYB2 (green), CWPs (red), and GalNAc enzymes (black) in response to encystation stimuli. GLP4 increases rapidly within 30 min and decreases at 4 h, MYB2 increases at 2-4h, CWPs increase at 2-4h, and enzymes in the GalNAc pathway increase by 7-18 h. ***Bottom panel*** shows the regulatory relationship of GLP4, MYB2, CWPs, and enzymes of the GalNac pathway in G1+S and G2+M cells after exposure to encystation stimuli. Bottom left indicates that GLP4 and MYB2 are not upregulated in the trophozoite (Pre-encystation). CWPs and GalNAc enzymes remain uninduced. Bottom right indicates that *Giardia* cells accumulate in G2+M phase in response to encystation stimuli (accumulation depicted by size of the blue arrows). MYB2 has been shown to bind the promoter regions of CWP1-3 and G6PI-B as well as function as an activator in previous studies. In G1+S phase, GLP4 is upregulated to repress MYB2, hence CWPs and GalNAc enzymes are downregulated. In G2+M phase, GLP4 is downregulated and MYB2 is upregulated. Therefore, CWPs and GalNAc enzymes are upregulated.

## Discussion

Previous studies revealed that TFs play important roles in coordinating the late stages of encystation. However, the contribution of TFs during the early stage of encystation was largely unknown. Here we systematically studied the expression profiles of 6 TFs in the early stage of encystation. We utilized CRISPRi- and CasRX-mediated knockdown tools to delineate the role of early upregulated TFs in encystation. We identified GLP4, a previously unstudied TF, as a cell cycle-dependent repressor of early encystation. GARP TFs are commonly considered to be plant-specific, but are also found in protists. Several plant GARP TFs are required for differentiation in *Arabidopsis*, such as reproductive organ determination (44). In our model, encystation stimuli cause *Giardia* trophozoites to accumulate at G2+M phase, and both G1+S and G2+M cells enter the encystation program (Fig. 5). Interestingly, GLP4 expression is rapidly upregulated in the G1+S population to repress MYB2 and CWP1 levels (Fig. 5). In contrast, GLP4 levels in the G2+M population is slightly decreased, thus allowing the activation of MYB2 to upregulate the expression of CWP1-2 as well as enzymes in the GalNAc pathway (Fig. 5). This model suggests that a lower dose of GLP4 in G2+M cells permits upregulation of MYB2 and encystation specific genes while a higher dose of GLP4 in G1+S cells downregulates MYB2 and the encystation program.

Our observations provide new insights into regulation of stage conversion in *Giardia*. First, our results indicate that GLP4 prevents some cells from encysting and therefore GLP4 has a role in controlling the balance between proliferative growth and terminal differentiation into infectious cysts. It would be interesting to extend our *in vitro* studies into an *in vivo* model of infection to confirm a role for GLP4 in regulating the balance between persistence of infection and transmission. Second, previous studies indicated negative cis-acting regulatory elements are present in CWP2 and G6PI-B promoters (32, 33). We have noticed CWP2 and G6PI-B promoters contain GARP binding sequence (A/G)ATCN (25). Although our results showed that CWP1-2, G6PI-B, UAE, and GNPNAT are upregulated in GLP4 mutants, how this newly identified repressor GLP4 interacts with promoter binding sequence and other TFs is worth further investigation. Third, phospho-regulation may play a key role in regulating entry into the encystation pathway. *Arabidopsis* nucleus-localized GARP TFs, type B ARRs, were activated through a phosphorylation cascade to modulate plant hormone cytokinin signaling (45). *Giardia’s* cyclin dependent kinase 2 (CDK2) is upregulated at G2+M (46) and has been shown to activate MYB2 through phosphorylation (47). Two recent RNAseq studies also identified an encystation-upregulated cyclin (GL50803_15532) (13) and G2-dependent TF GLP1 (GL50803_7272) (46). In the future, it would be interesting to explore how GLP1, GLP4, CDK2, and cyclin work together to initiate encystation.

Our findings advance the understanding of the dynamics of TFs expression in the early phase of encystation in *Giardia*. Moreover, our results demonstrate an unexpected role for the cell cycle-dependent GLP4 transcriptional repressor in regulating MYB2 and the cyst production pathways. The theory of dosage-dependent transcriptional responses is essential for understanding how mammalian and plant cells modulate differentiation (48). Our findings on MYB2 and GLP4 suggests that *Giardia*, a single-celled organism, employs dosage-dependent transcriptional responses to tune the initiation of differentiation.

## Methods

### Giardia growth and encystation media

*Giardia intestinalis* isolate WB clone C6 (ATCC catalog number 50803; American Type culture collection) were cultured in TYDK media at pH 7.1 supplemented with 10% adult bovine serum and 0.125 mg/ml bovine bile(49). To induce encystation, cells were cultured 48 h in pH 6.8 pre-encystation media without bile then one of three encystation protocols were used: (1) Uppsala encystation protocol: TYDK media at pH 7.8 supplemented with 10% adult bovine serum and 5 mg/ml bovine bile(14); (2) Two-step protocol: TYDK media at pH 7.8 supplemented with 10% adult bovine serum, 0.25 mg/ml porcine bile and 5 mM lactic acid(50); (3) Lipoprotein-deficient protocol: TYDK media at pH 7.8 supplemented with lipoprotein-deficient serum(40).

### Plasmid construction

#### mNeonGreen and NanoLuc fusions

Coding sequences were PCR-amplified from *Giardia lamblia* genomic DNA. Primers sequences are indicated in the supplemental excel file Table S1. The mNeonGreen and NanoLuc vectors were digested with the indicated restriction enzymes and a PCR amplicon was ligated using Gibson assembly(51). The resulting constructs were linearized with the restriction enzyme indicated in supplemental Table S1 before electroporation for integration into the endogenous locus(52). Neomycin and puromycin were used for selection.

#### CasRX expression cassette design

EsCas13d (Catalog #108303), CasRX (Cat #109049), and PspCas13b (Cat#103862) were obtained from Addgene. Cas 13 fragments were PCR amplified, digested with restriction enzymes and inserted into pPAC-3HA expression cassettes under the GDH promoter(53). The Cas13-3HA expression vectors were linearized with SwaI and electroporated into *Giardia lamblia* so that expression levels could be monitored by western blotting. To generate the CasRX expression system, the CRISPRi expression vector (dCas9g1pac) was used as a backbone and dCas9 was replaced with CasRX and the gRNA scaffold sequence (SCF) was replaced with the CasRX-specific direct repeat (DR) (37).

#### Design of guide RNA

Guide RNA for the CRISPRi system utilized the Dawson Lab protocol (34), NGG PAM sequence and G. Lamblia ATCC 50803 genome were selected for CRISPRi guide RNA design with Benchling. Cas13 guide RNA designs were based on the Sanjana Lab Cas13 guide tool (https://cas13design.nygenome.org/)(54).

#### In vitro bioluminenscence assays

*Giardia* cells were iced for 15 min and centrifuged at 700 x g for 7 min at 4°C. Cells were resuspended in cold 1X HBS (HEPES-buffered saline) and serial dilutions were made after counting cells with a MOXI Z mini Automated Cell Counter Kit (Orflo, Kenchum, ID). To measure NanoLuc luminescence, 20,000 cells were loaded into white polystyrene, flat bottom 96-well plates (Corning Incorporated, Kennebunk, ME) then mixed with 10 μl of NanoGlo luciferase assay reagent (Promega). Relative luminescence units (RLU) were detected on a pre-warmed 37°C EnVision plate reader (Perkin Elmer, Waltham, MA) for 30 min to reach the maximum value. Experiments are from three independent bioreplicates. To measure CBG99 luminescence, 20,000 cells were loaded into white polystyrene, flat bottom 96-well plates then mixed with 50 μl of 10 mg/mL D-luciferin. Relative luminescence units (RLU) were detected as above.

#### Protein blotting

*Giardia* parasites were iced for 30 min then centrifuged at 700 x g for 7 min and washed twice in 1X HBS supplemented with HALT protease inhibitor (Pierce) and phenylmethylsulfonyl fluoride (PMSF). The cells were resuspended in 300 μl of lysis buffer containing 50 mM Tris-HCl pH 7.5, 150 mM NaCl, 7.5% glycerol, 0.25 mM CaCl_2_, 0.25 mM ATP, 0.5 mM Dithiotheitol, 0.5 mM PMSF (Phenylmethylsulfonyl flroride), 0.1% Trition X-100 and Halt protease inhibitors (Pierce). The sample was pelleted at 700 x g for 7 min, the resulting supernatant was mixed with 2X sample buffer (Bio-Rad) and boiled at 98°C for 5 min. Protein samples were separated using sodium dodecyl sulfate (SDS) polyacrylamide gel electrophoresis. Protein transfer was performed using an Immobilon-FL membrane (Millipore). To detect tubulin, a mouse monoclonal anti-acetylated tubulin clone 6-11B-1 antibody (IgG2b; product T 6793; Sigma-Aldrich) were used at 1:2,500 dilution and secondary anti-mouse isotype-specific antibody conjugated with Alexa 488 (anti-IgG2b) were used at 1:2,500. To detect CWP1, Alexa 647-conjugated anti-CWP1 antibody (Waterborne, New Orleans, LA) was used at 1:2,000. Multiplex immunoblots were imaged using a Chemidoc MP system (Bio-Rad).

#### Immunofluorescence

*Gairdia* parasites were iced for 30 min and pelleted at 700 x g for 7 min. The pellet was fixed in PME buffer (100 mM Piperazine-N,N’-bis (ethanesulfonic acid) (PIPES) pH 7.0, 5 mM EGTA, 10 mM MgSO_4_ supplemented with 1% paraformaldehyde (PFA) (Electron Microscopy Sciences, Hatfield, PA), 100 μM 3-maleimidobenzoic acid N-hydroxysuccinimide ester (Sigma-Aldrich), 100 μM ethylene glycol bis (succinimidyl succinate) (Pierce), and 0.025% Triton X-100 for 30 min at 37°C. Fixed cells were attached to polylysine coated coverslips. Cells were washed once in PME and permeabilized with PME plus 0.1% Triton X-100 for 10 min. After two quick washes with PME, blocking was performed in PME supplemented with 1% bovine serum albumin, 0.1% NaN_3_, 100 mM lysine, 0.5% cold water fish skin gelatin (Sigma-Aldrich). Next, 1:200 diluted Alexa 647-conjugated anti-CWP1 antibody (Waterborne, New Orleans, LA) was added to incubate for 1 h. Cells were washed three times in PME plus 0.05% Triton X-100. Coverslips were mounted with ProLong Gold antifade plus 4’, 6-diamidino-2-phenylinodole (DAPI; Molecular Probes). Images were acquired on a DeltaVision Elite microscope using a 60X, 1.4-numerical aperture objective with a PCO Edge sCMOS camera, and images were deconvolved using SoftWorx (API, Issaquah, WA).

#### Imaging and image analysis

Analyses of CWP1-stained vesicle size and number were performed with Imaris software (Bitplane, version 8.1). ImageJ was used to process all images and figures were assembled using Adobe Illustrator.

#### Cyst count and cyst viability staining

To collect water-resistant *Giardia* cysts, confluent *Giardia* trophozoites were incubated in encystation media supplemented with 10 g/L ovine bovine bile and calcium lactate. To count cyst, 20 μl of 48 h encysted cells were counted using a hemocytometer. To determine cyst viability, 48 h encysted *Giardia* cells were centrifuged at 700 x g for 7 min and the pellets were washed 10 times in deionized water, then stored in distill water overnight at 4°C. The next day, fluorescein diacetate (FDA) and propidium iodide (PI) were used to stain live and dead cysts, respectively, and images were collected using a DeltaVision Elite microscope with a 40X, 1.4-numerical apeture objective with a PCO Edge sCMOS camera, and images were deconvolved using SoftWorx (API, Issaquah, WA).

#### Flow cytometry assay

Flow cytometry analysis was performed after fixation with 0.25% PFA at 4°C for 15 min. 1 μM DRAQ5 (Thermo Scientific Cat# 62251) was used to stain DNA. 10,000 cells per sample were analyzed using a FACS Canto II Flow Cytometer at the Pathology Flow Cytometry Core Facility (Department of Pathology, University of Washington). Data were analyzed using FlowJo.

## Acknowledgements

We thank Paredez laboratory undergraduate students Bailin Zhang, Greyson Hamilton, and Daria Rydell and our colleagues at UW Biology for discussions and comments on the manuscripts, and Xiao Ping from the flow cytometry facility at UW Pathology. We thank Photini Sinnis, Kirk Deitsch, Patricia Johnson for inviting us to participate in the 40^th^-Biology of Parasitism (BoP) course where BoP students helped to initiate this project. We thank Dr. Scott Dawson for gifting us the CRISPRi plasmid and Dr. Barbara Wakimoto and Dr. Melissa Steele-Ogus for providing comments on the manuscript. Research was funded by University of Washington Bridge funds and NIAD 1R21AI159035 to ARP.

---

**Fig. S1.**
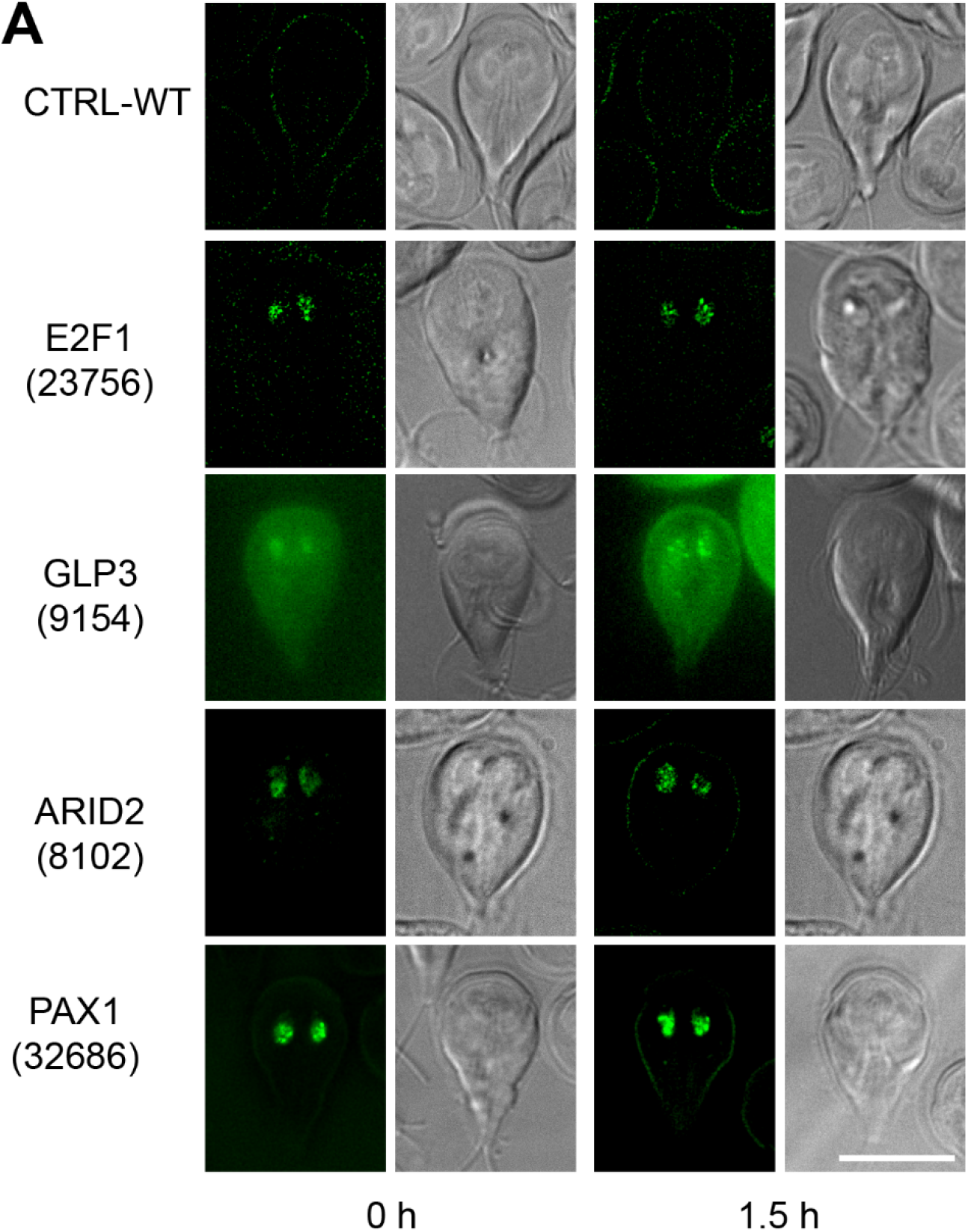
Localizations of encystation-induced genes. mNG and DIC images of Wild Type (untagged control cell), and mNG-tagged proteins at the endogenous locus: E2F1 (GL50803_23756), GLP3 (GL50803_9154), ARID2 (GL50803_8102), PAX1 (GL50803_32686) after 0 and 1.5 h exposure to encystation medium. Bars, 10 μm.

**Fig. S2.**
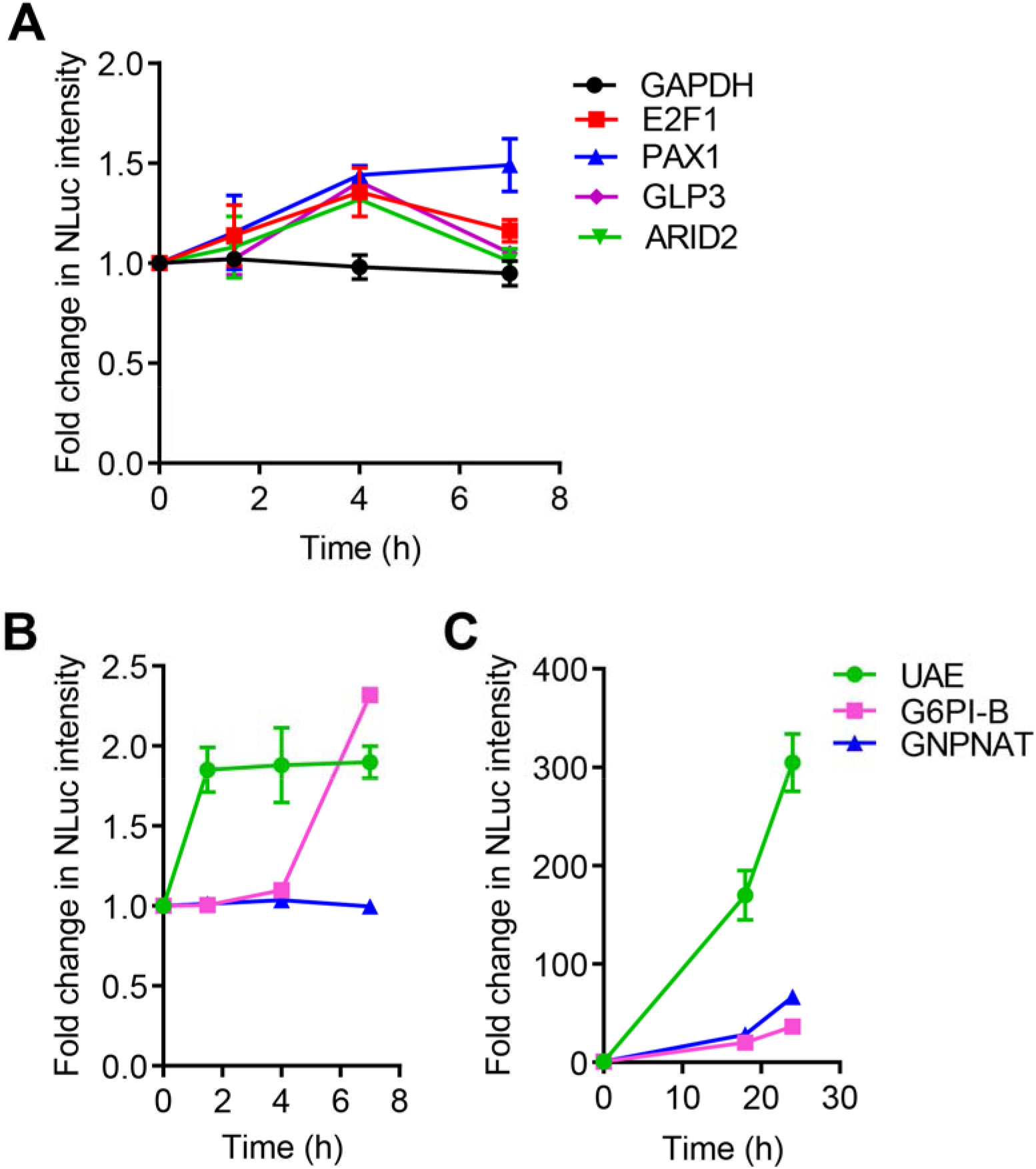
E2F1, PAX1, GLP3, and ARID2 are upregulated at 4h. *(A)* Relative expression of NanoLuc intensity for E2F1-NLuc, PAX1-NLuc, ARID2-NLuc, GLP3-NLuc, and GAPDH-NLuc after 0, 1.5, 4, and 7 h exposure to encystation medium. The expression level from each time point has three biological replicates. The fold change is normalized to the 0 h control. *(B-C)* G6PI-B (GL50803_8245), UAE (GL50803_7982), and GNPNAT (GL50803_14259) are upregulated in response to encystation stimuli. Relative NanoLuc intensity of G6PI-B, UAE, and GNPNAT were measured after 0, 1.5, 4, 7, 18, and 24 h exposure to encystation medium. The expression level from each time point has three biological replicates. The fold change is normalized to 0 h.

**Fig. S3.**
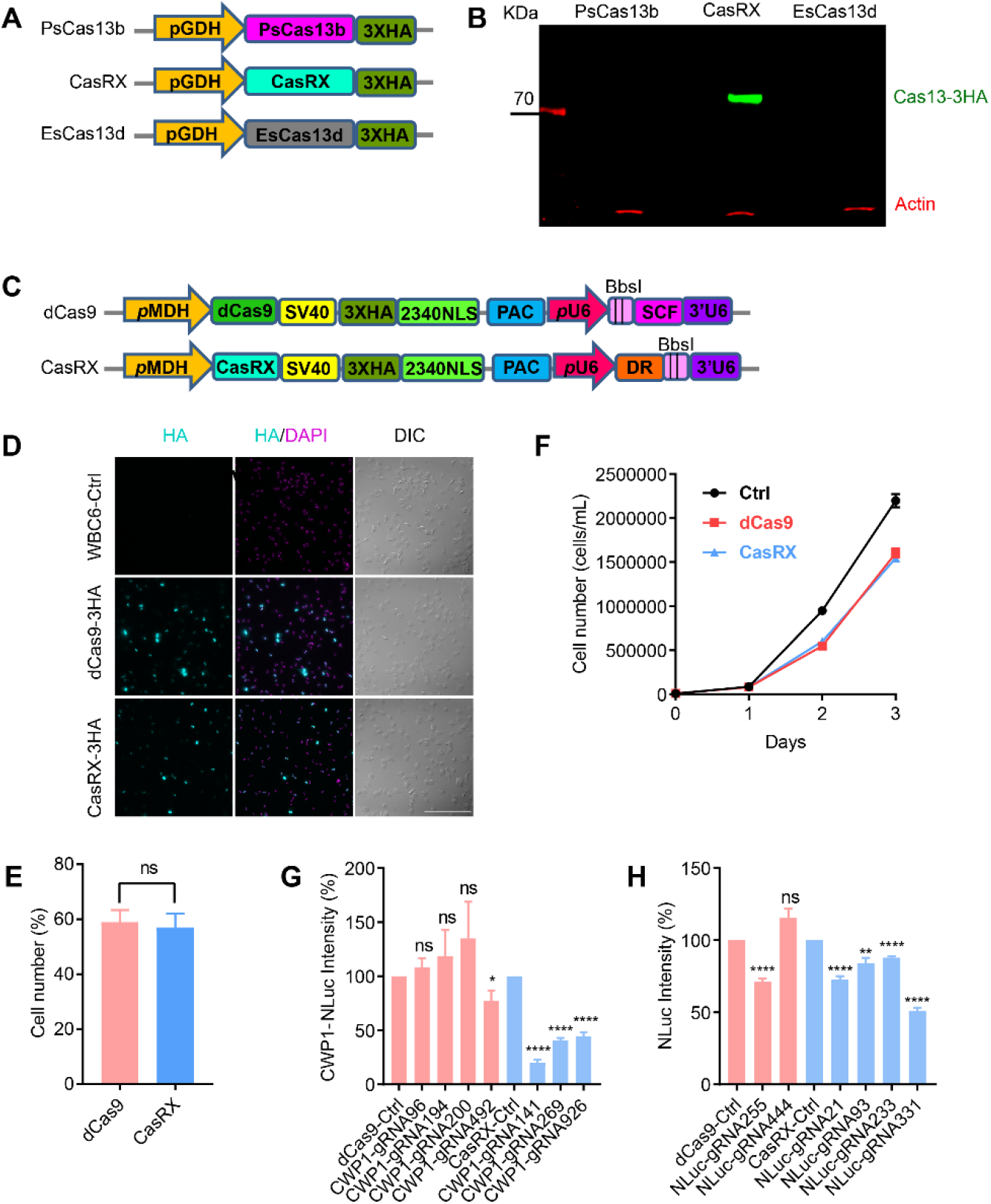
Development of CasRX for use in *Giardia*. Cas13 enzymes are RNA-targeting CRISPR enzymes which have been shown to have high efficiency and specificity in human cells (37). We first tested the ability to express three Cas13 enzymes, including PsCas13b (55), CasRX (37) and EsCas13d (37). CasRX was the only Cas13 enzyme with good expression in *Giardia*. Starting with the existing CRISPRi vector, we replaced dCas9 and the gRNA scaffold sequence (SCF) with CasRX and the CasRX-specific direct repeat (DR), to establish a functional CasRX knockdown system. *(A,B)* Heterologous expression of 3HA tagged PsCas13b, CasRX, and EsCas13d in *Giardia*. pGDH= promoter of glyceraldehye 3-phosphate dehydrogenease (GL50803_6877). HA=Hemagglutinin-tag. *(C)* Schematic of CRISPRi vector dCas9g1pac and CasRX vector design. pMDH= promoter of malate dehydrogenase, pac=puromycin resistance marker, 2340NLS=GL50803_2340 nuclear localization signal, SCF=gRNA scaffold sequence, DR=Direct repeat for CasRX, *pU6=Giardia* U6 promoter. To determine if one of these knockdown systems might be better tolerated than the other, we examined the proportion of cells that expressed dCas9 or CasRX and found no appreciable difference. *(D)* Localizations of 3XHA tagged dCas9 and CasRX with DAPI staining in *Giardia. (E)* Quantification of cells with detectable dCas9 and CasRX expression. n.s.=not significant (p=0.70). Data are mean ± s.d. from three independent experiments (dCas9 n=3278, CasRX n=3327 cells) Mann-Whitney U-test. *(F)* Growth curve of dCas9 and CasRX expressing cell lines indicates that the growth rate is essentially identical for dCas9 and CasRX expressing cells. Blue=CasRX, Red=dCas9. Bars, 100 μm. To compare efficacy of knockdown, we designed CasRX- and CRISPRi-specific guide RNAs to deplete NanoLuc and CWP1. For these two genes CasRX outperformed CRISPRi reaching up to 80% reduction for CWP1. *(G-H)* Relative NanoLuc intensity of proteins targeted for knockdown with CRISPRi or CasRX gRNAs. Levels were normalized to non-specific gRNA controls. *(G)* Quantification of NanoLuc intensity of CWP1 (GL50803_5638), and *(H)* NanoLuc only. The expression levels is normalized to guide RNA controls. All experiments are from three independent biological replicates. Data are mean ± s.d. (n=3). Student’s t-test, *p<0.01, **p<0.001, ***p<0.0001, ****p<0.00001, ns= not significant.

**Fig. S4.**
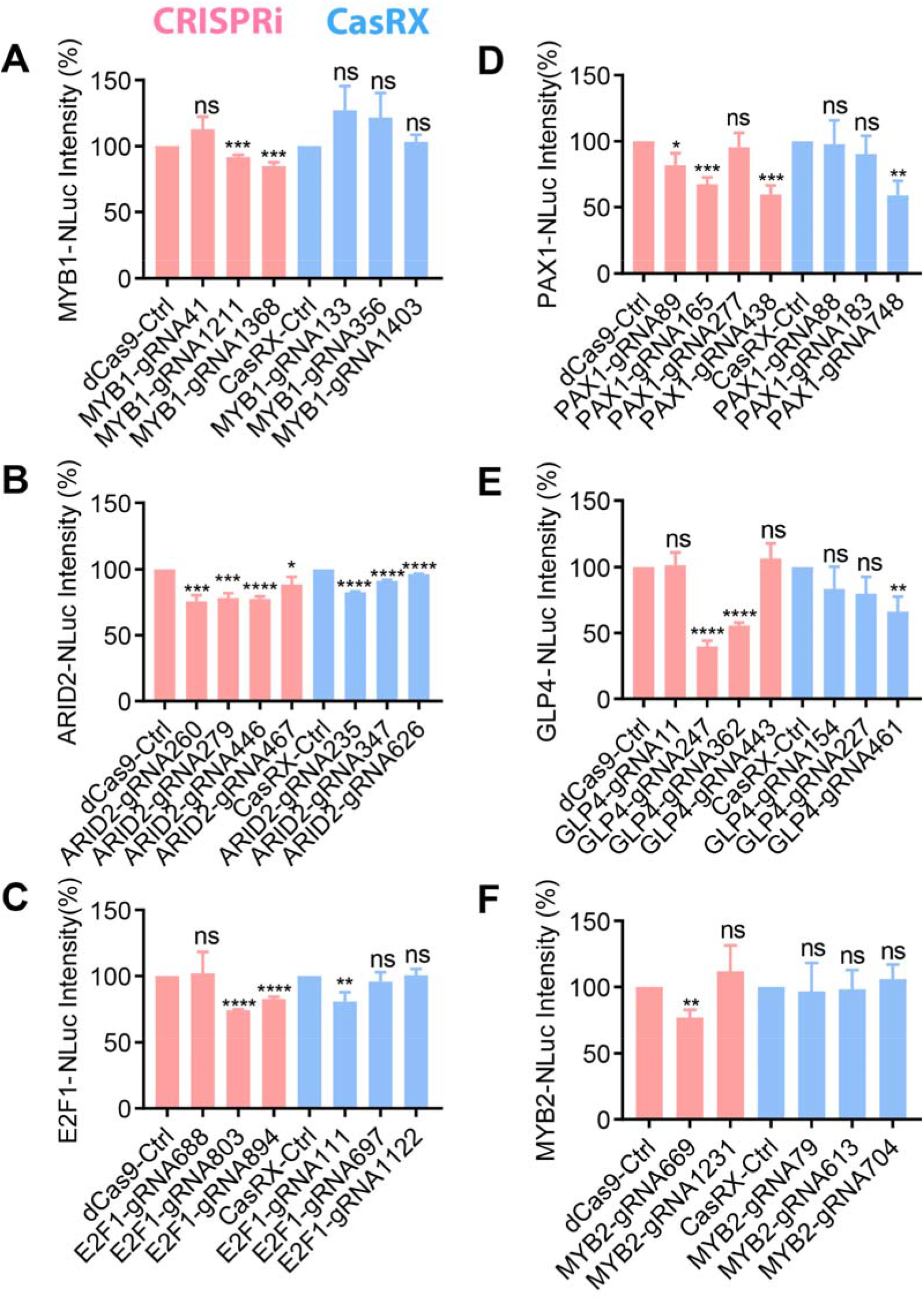
Screen of guide RNAs with CRISPRi and CasRX systems. *(A-F)* Quantification of NanoLuc intensity using the indicated CRISPRi or CasRX guide RNAs. Levels were normalized to non-specific gRNA controls. *(A)* MYB1-NLuc (GL50803_5347). *(B)* ARID2-NLuc (GL50803_8102). *(C)* E2F1-NLuc (GL50803_23756). *(D)* PAX1-NLuc (GL50803_32686). *(E)* GLP4-NLuc (GL50803_33232). *(F)* MYB2-NLuc (GL50803_8722). All experiments are from three independent biological replicates. Data are mean ± s.d. (n=3) Student’s t-test, *p<0.01, **p<0.001, ***p<0.0001, ****p<0.00001, ns= not significant.

**Fig. S5.**
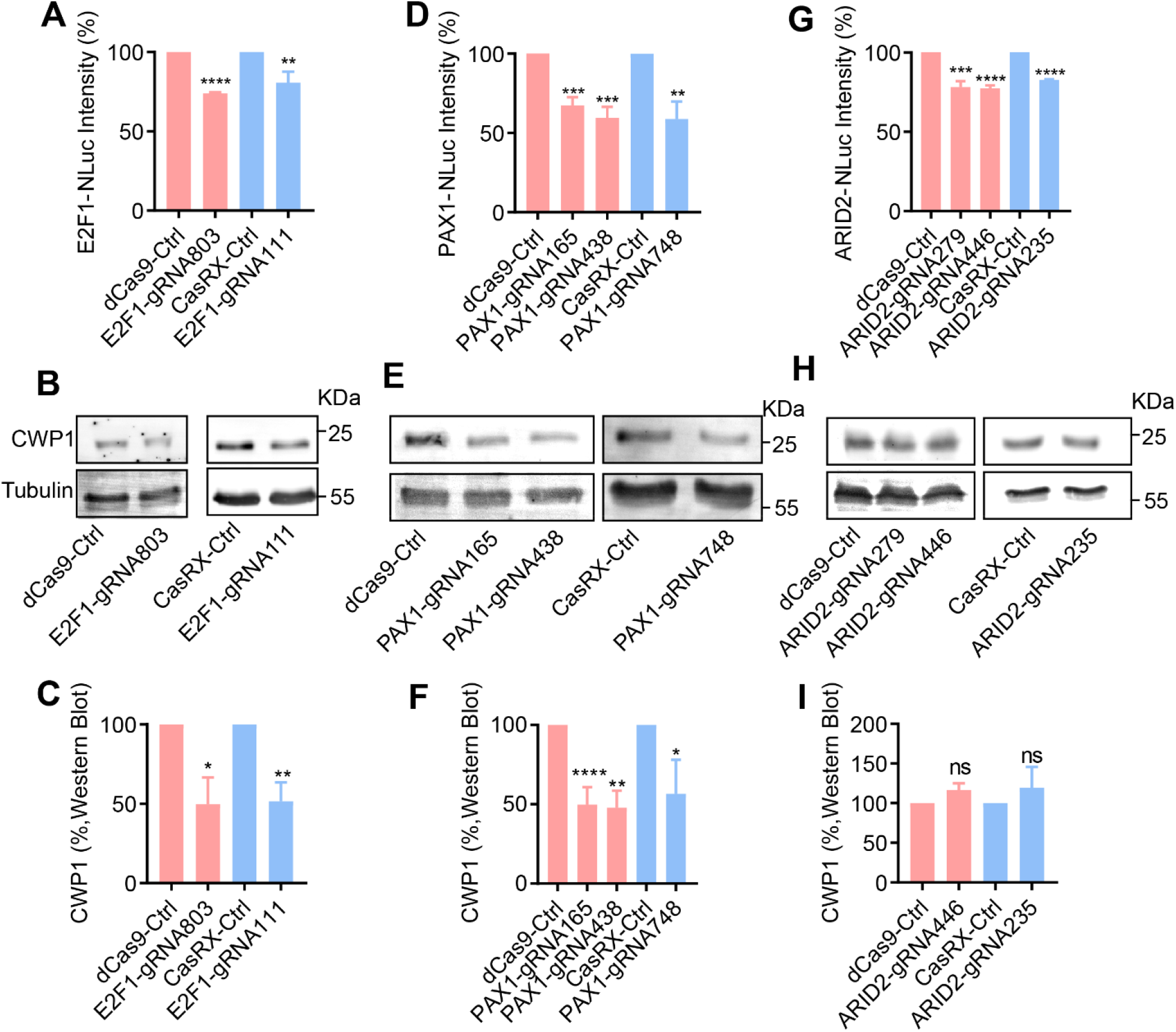
Knockdowns of E2F1, PAX1, and ARID2 in *Giardia. (A,D,G)* Quantification of NanoLuc intensity from CRISPRi (or CasRX)-mediated knockdowns of (A) E2F1 (GL50803_23756), (D) PAX1 (GL50803_32686), and (G) ARID2 (GL50803_8102). The expression level is normalized to the corresponding dCas9 or CasRX control. *(B,E,H)* Western blots of CWP1 and tubulin from (B) E2F1, (E) PAX1, and (H) ARID2 knockdowns at 4 h of encystation. *(C,F,I)* Quantification of Western blots from (B) E2F1, (E) PAX1, and (H) ARID2 knockdowns, respectively. The expression level is normalized to the tubulin control. All experiments are from three independent biological replicates. Data are mean ± s.d. (n=3) Student’s t-test, *p<0.01, **p<0.001, ***p<0.0001, ****p<0.00001, ns= not significant.

**Fig. S6.**
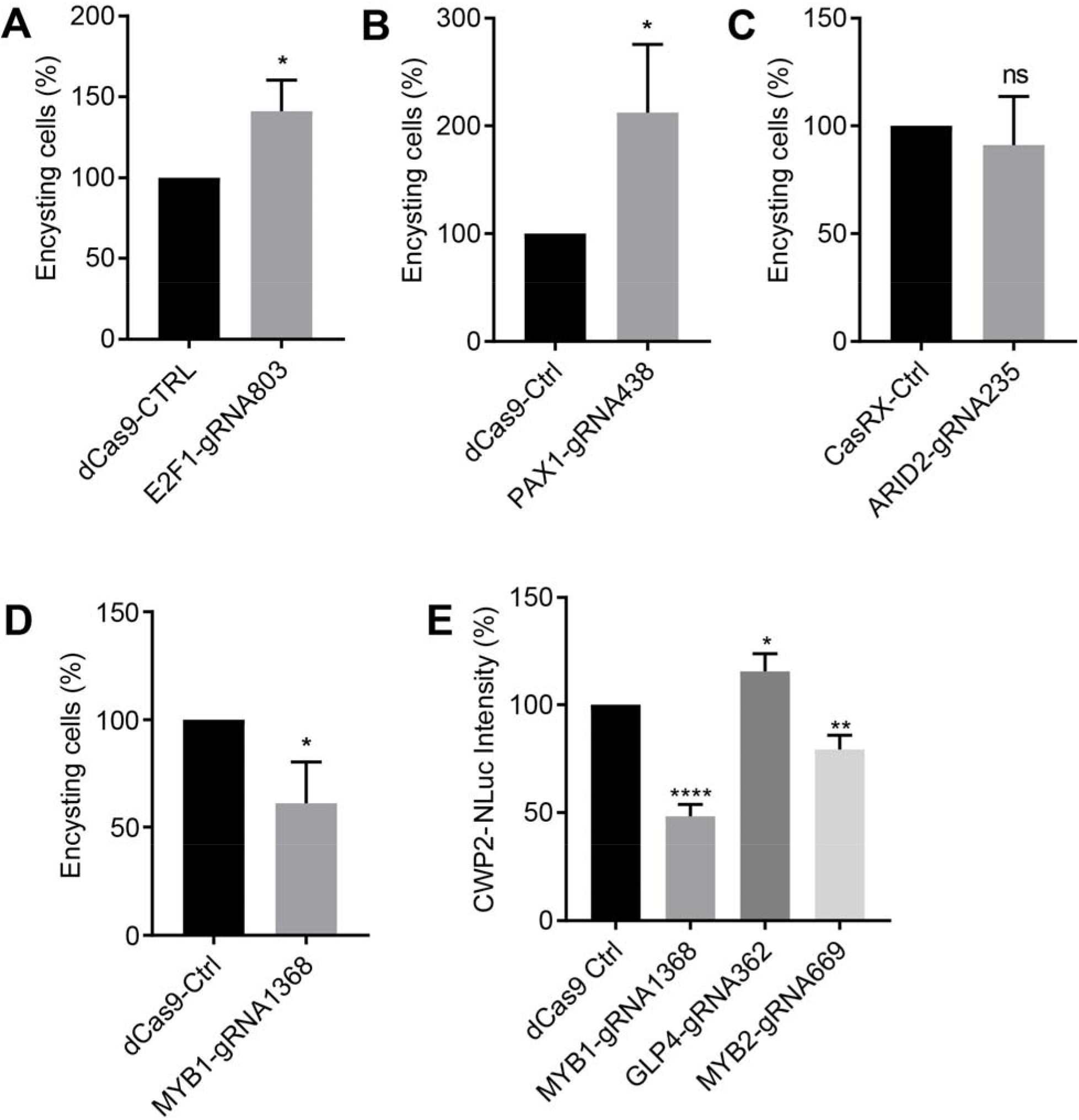
Proportion of encysting cells in knockdowns of (A) E2F1, (B) PAX1, (C) ARID2, and (D) MYB1 at 4 h post induction of encystation. Data are mean± s.d. from three independent experiments (total cells counted for ***A***, dCas9-Ctrl n=626, E2F1-gRNA803 n=655; ***B***, dCas9-Ctrl n=638, PAX1-gRNA438 n=640; ***C***, CasRX-Ctrl n=625, ARID2-gRNA235 n=447 cells, **D**, dCas9-Ctrl n=428, MYB1-gRNA1368 n=356) Student’s t-test, *p<0.01, **p<0.001, ****p<0.00001, ns= not significant. *(E)* Relative expression of CWP2 NanoLuc intensity in knockdowns of MYB1, MYB2 and GLP4 at 4 h post induction of encystation. The expression level is normalized to the dCas9 Ctrl. All experiments are from three independent biological replicates. Data are mean ± s.d. (n=3) Student’s t-test, *p<0.01, **p<0.001, ****p<0.00001.

**Fig. S7.**
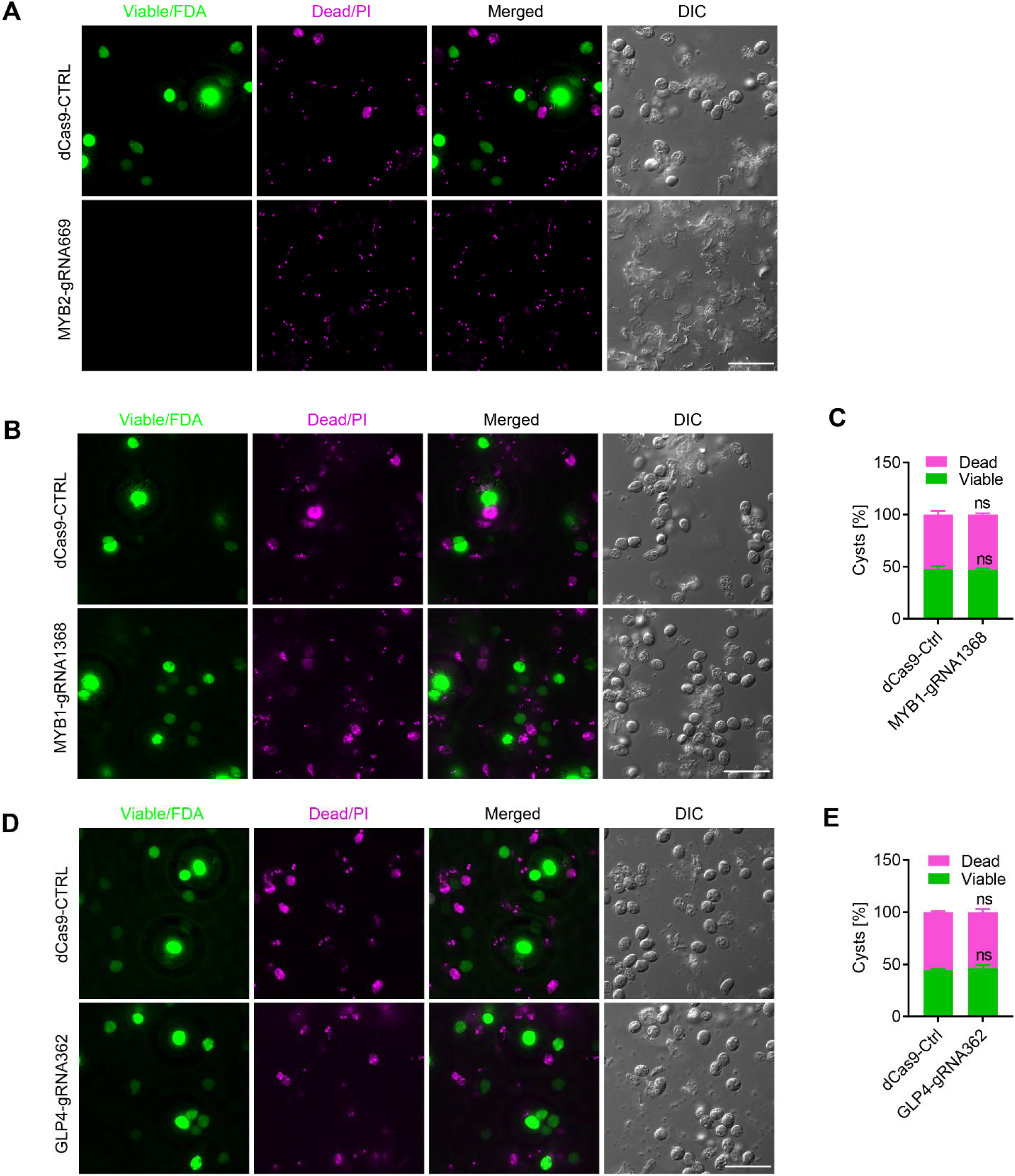
Knockdown of GLP4 and MYB1 do not alter cyst viability. *(A)* MYB2-gRNA669 derived cysts stained with FDA (fluorescein diacetate, green=live) and PI (propidium iodine, magenta=dead) to determine cyst viability. *(B)* MYB1-gRNA1368 and (D) GLP4-gRNA362 derived cysts stained with FDA and PI. *(C)* Quantification of *(B). (E)* Quantification of (D). All cell lines were encysted for 48 h. Data are mean ± s.d. from three biological replicates (total cysts counted for dCas9-Ctrl n=400, GLP4-gRNA362 n=499, MYB1-gRNA1368 n=399) Student’s t-test, ns= not significant. Bars, 100 μm.

**Fig. S8.**
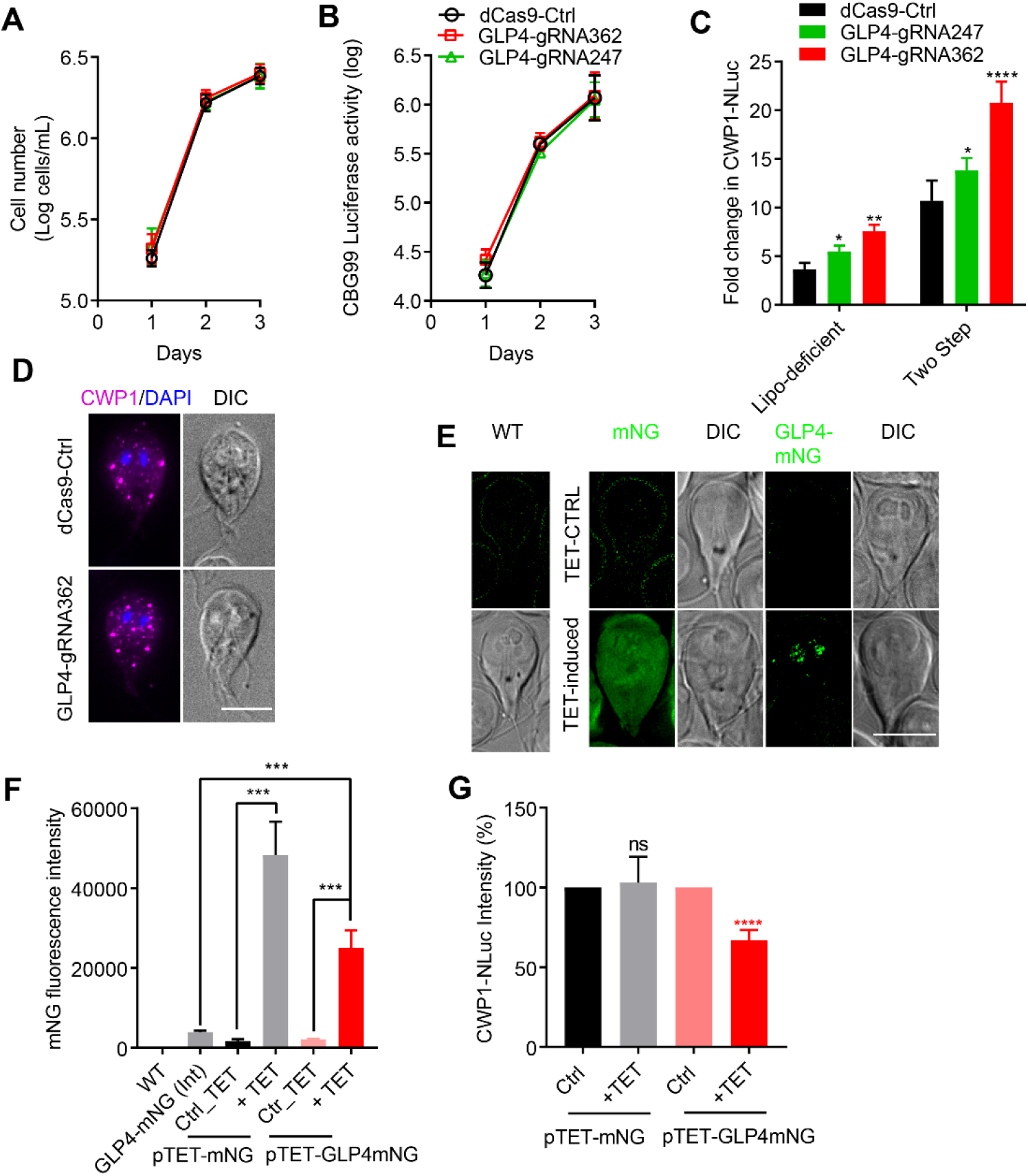
Knockdown of GLP4 does not alter growth rate or cell viability. *(A)* Cell growth assay for dCas9 control and GLP4 knockdown strains. All cultures were started with 200,000 cells and then cell concentration was determined with a MOXI Z counter at 1, 2, and 3 days. *(B)* Cell viability of dCas9 control and GLP4 knockdowns at 1, 2, 3 days as determined by ATP dependent luminescence from CBG99. Three independent replicates of 200,000 cells/well were assayed in a plate reader. Black, dCas9 control. Red, GLP4-gRNA362. Green, GLP4-gRNA247. *(C)* Relative CWP1 NanoLuc intensity in GLP4 knockdowns at 4 h of encystation with lipoprotein-deficient medium or two step encystation medium. *(D)* Representative images of CWP1 stained vesicles in dCas9 control and GLP4-gRNA362 cell lines. *(E-F)* mNG expression images of WT, GLP4-mNG (Int) (integrated and endogenously tagged GLP4-mNG), pTET-mNG (mNG under the tetracycline inducible promoter) and pTET-GLP4-mNG (GLP4-mNG under the tetracycline inducible promoter) in the CWP1-NLuc background with or without tetracycline induction. All images were taken with same exposure. Bars, 10 μm. 20 μg/ml tetracycline (final concentration) was added to induce expression. *(F)* Quantification of (E). 2x 10^6^ cells were assayed in each well of a clear-bottom black plate and mNG fluorescence intensity detected with a fluorescent plate reader. *(G)* Relative CWP1-NLuc intensity in mNG and GLP4-mNG overexpression cell lines at 4 h of encystation. Data are mean ± s.d. (n=3) Student’s t-test, *p<0.01, **p<0.001, ****p<0.00001, ns=not significant.

**Fig. S9.**
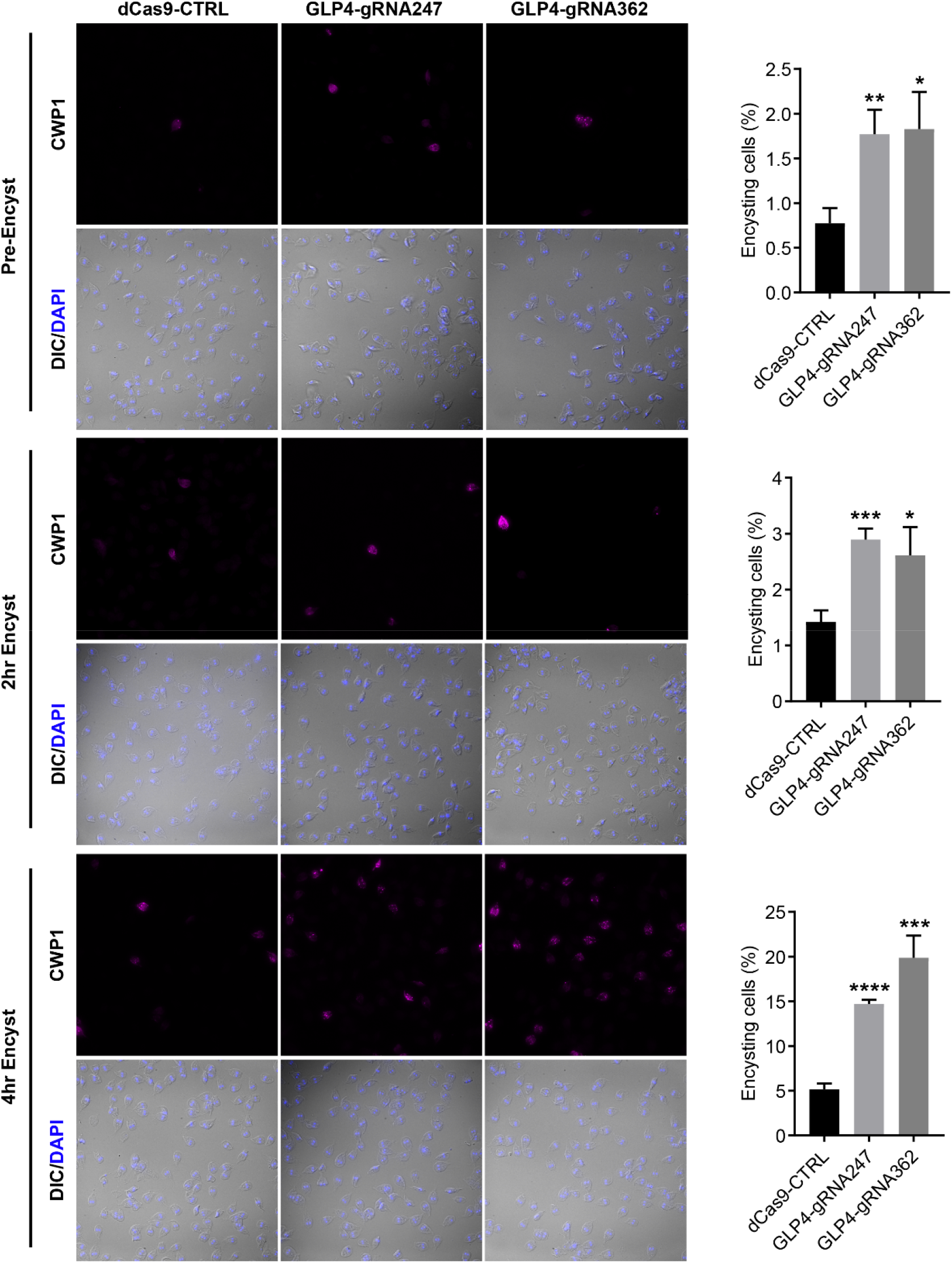
Knockdown of GLP4 increases encystation rates. Quantification of CWP1 positives cells in control and GLP4 knockdown strains at 0 h, 2h, and 4h post induction of encystation. Data are mean ± s.d. from three biological replicates (total cells counted for 0 h: dCas9-Ctrl n=4927, GLP4-gRNA247 n=4927, GLP4-gRNA362 n=5808; 2h: dCas9-Ctrl n=5785, GLP4-gRNA247 n=5785, GLP4-gRNA362 n=6041, 4 h: dCas9-Ctrl n=5312, GLP4-gRNA247 n=5312, GLP4-gRNA362 n=4988 cells) Student’s t-test, *p<0.01, **p<0.001, ***p<0.0001, ****p<0.00001. Bars, 100 μm.

**Fig. S10.**
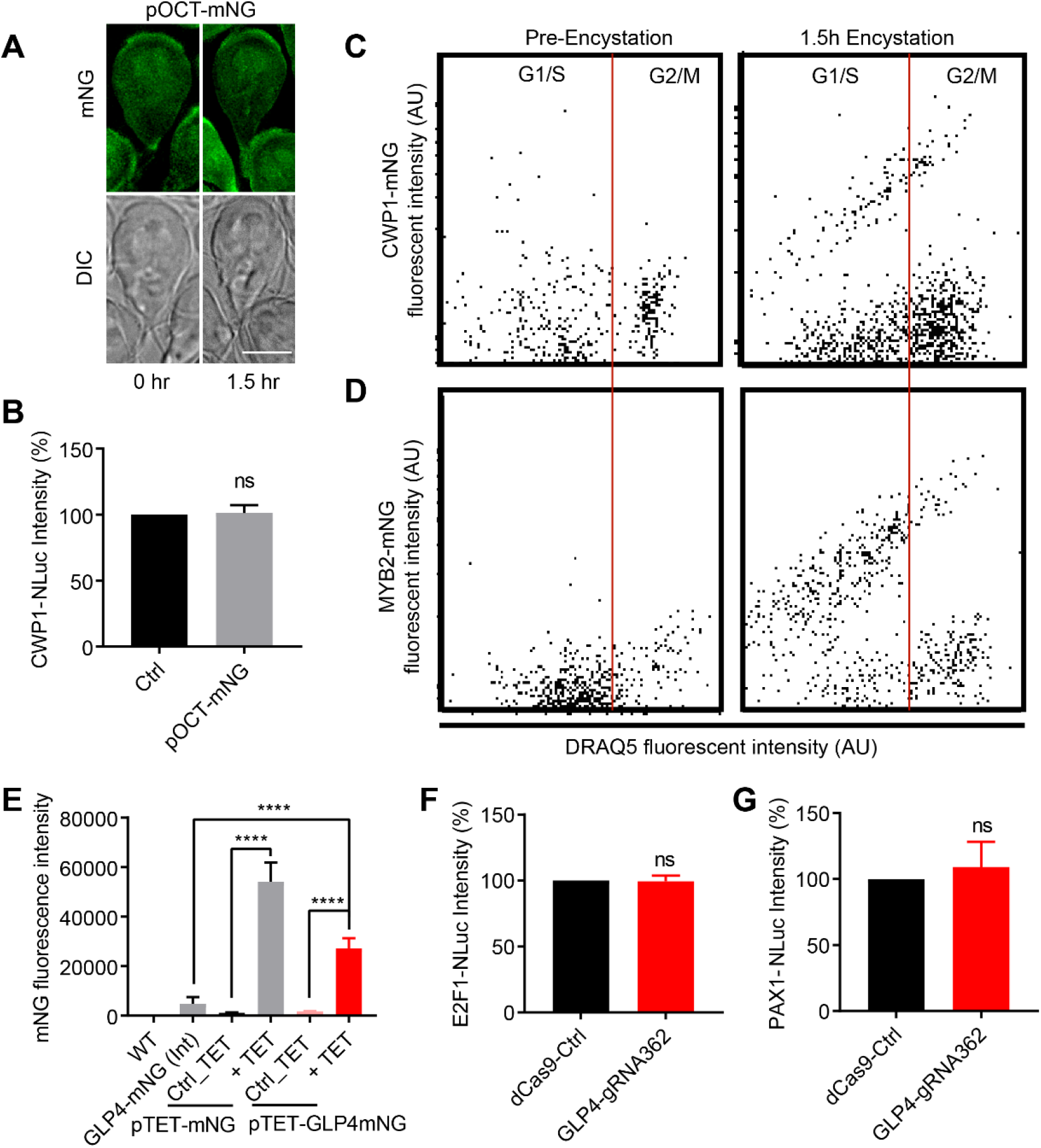
G2+M cells have higher levels of CWP1 and MYB2. *(A)* Fluorescent images of the pOCT::mNG expressing cell lines at 1.5h post induction of encystation. pOCT=promoter of ornithine carbamoyltransferase (GL50803_10311), Bars, 10 μm. *(B)* Relative CWP1-NLuc intensity from pOCT-mNG cell lines at 4 h post induction of encystation. CWP1-NLuc intensity is normalized to the CWP1-NLuc parental line. Data are mean ± s.d. (n=3) Student’s t-test, n.s.=not significant. *(C)* CWP1-mNG fluorescence intensity of DRAQ5-stained cells at 0 and 1.5 h of encystation. *(D)* MYB2-mNG fluorescence intensity of DRAQ5-stained cells at 0 and 1.5 h encystation. *(E)* mNG fluorescent intensity of WT, GLP4-mNG (Int) (integrated and endogenously tagged GLP4-mNG), pTET-mNG (mNG under the tetracycline inducible promoter) and pTET-GLP4-mNG (GLP4-mNG under the tetracycline inducible promoter) in the MYB2-NLuc cells with or without tetracycline induction. 20 μg/ml tetracycline (final concentration) was added to induce expression. Relative NanoLuc intensity of *(F)* E2F1 and *(G)* PAX1 levels in GLP4 knockdown cell lines at 4 h of encystation. Expression levels are normalized to the dCas9 control at 4h post induction of encystation. Bars, 10 μm. Data are mean ± s.d. (n=3) Student’s t-test, *p<0.01, **p<0.001, ****p<0.00001, n.s.=not significant.

Table S1 Supplemental excel file containing all of the primer sequences used in this study.

